# Functionalising the electrical properties of Kombucha zoogleal mats for biosensing applications

**DOI:** 10.1101/2024.01.21.576528

**Authors:** Anna Nikolaidou, Alessandro Chiolerio, Mohammad Mahdi Dehshibi, Andrew Adamatzky

## Abstract

Kombucha is a type of tea that is fermented using yeast and bacteria. During this process, a film made of cellulose is produced. This film has unique properties such as biodegradability, flexibility, shape conformability, and ability to self-grow, as well as be produced across customised scales. In our previous studies, we demonstrated that Kombucha mats exhibit electrical activity represented by spikes of electrical potential. We propose using microbial fermentation as a method for *in situ* functionalisation to modulate the electroactive nature of Kombucha cellulose mats, where graphene and zeolite were used for the functionalisation. We subjected the pure and functionalised Kombucha mats to mechanical stimulation by applying different weights and geometries. Our experiments demonstrated that Kombucha mats functionalised with graphene and zeolite exhibit memfractive properties and respond to load by producing distinctive spiking patterns. Our findings present incredible opportunities for the *in situ* development of functionalised hybrid materials with sensing, computing, and memory capabilities. These materials can self-assemble and self-grow after fusing their living and synthetic components. This study contributes to an emergent area of research on bioelectronic sensing and hybrid living materials, opening up exciting opportunities for use in smart wearables, diagnostics, health monitoring and energy harvesting applications.

## 1. Introduction

There has been a significant increase in the use of cellulose as an alternative sustainable and renewable material in various fields like medicine, packaging, food, and textiles. Cellulose is a natural biodegradable polymer and is considered one of the most abundant macromolecules in Nature. It is produced by a wide range of living organisms, from bacteria to plants [1, 2, 3]. Bacterial cellulose is known to exhibit higher purity, mechanical strength, water-holding capacity, chemical stability, and biological adaptability than cellulose produced by plants. This is due to a lack of lignin and hemicellulose polymers [4]. Acetic acid bacteria produce acetic acid by metabolising carbon sources into acetic acid and ethanol through their oxidative fermentation pathways [5]. They are excellent producers of cellulose [6], which can be obtained in a very short time [7, 8]. Cellulose is organised in hydrogels and features interesting trains of spikes of extracellular electrical potential [9]. These potential spikes could be used to enable reactive sensing wearables by means of living colonies of bacteria [10]. When a community of bacteria, such as acetic acid bacteria and yeast, form a symbiotic relationship, they can enhance cellulose production through their combined microbial metabolism [4]. This synergy is found in Kombucha cellulose, which emerges during the fermentation of Kombucha tea. Kombucha tea is a popular probiotic beverage that is made by fermenting sugared tea with a symbiotic culture of over 20 species of bacteria and yeast, known as SCOBY [11, 12, 13]. During fermentation, the microorganisms consume sucrose as the primary carbon source, while the tea extract provides the nitrogen source. In the presence of oxygen, SCOBY produces organic acids, carbon dioxide and a cellulosic floating biofilm composed of cellulose. The thickness of the film varies depending on the nutrients and breeding time [14]. Kombucha cellulose shows high biocompatibility due to its enhanced cell-matrix interactions, cell signalling pathways and ability to maintain cellular homeostasis [15]. In our previous studies, we have demonstrated that Kombucha cellulose mats show rich dynamics of electrical activity, represented by spikes of electrical potential. This makes them an exciting material for developing living bioelectronic materials with active electrical properties, sensing and computing capabilities [16, 17]. The electroactive nature of the cellulose mats also makes them suitable for biosensing applications.

The live Kombucha cellulose mats possess various chemical, electrical, and mechanical properties. They have a high production rate of cellulose, are cost-effective, flexible, and can take customised shapes while retaining the ability to self-grow. These mats can be produced in different sizes and thicknesses, making them an ideal material for scaffold production. They can be used to create biocomposites with targeted and enhanced functionalities, suitable for applications such as smart wearables, soft robotics, and building materials.

To enhance the electrical properties and sensing capabilities of Kombucha cellulose mats, in this study we combine graphene and zeolite for the functionalisation of the cellulose mats. By taking the direction of combining two materials that have distinct but collaborative properties [18, 19, 20], our hypothesis is that we can obtain a fabricated biocomposite material with higher sensitivity, stability and tunable electrochemical responses towards exposure to mechanical pressure. Graphene exhibits excellent electrical conductivity, mechanical flexibility, thermal conductivity, and low coefficient of thermal expansion behaviour [21, 22, 23, 23, 24, 25] and has been used as an alternative carbon-based nanofiller in the preparation of polymer nanocomposites [26, 27, 28, 29, 30, 31, 32, 33], for the fabrication of nano-electronic and bio-electronic devices [34] and to replace metal conductors in electronic and electrical devices [35, 36, 37]. Zeolites are crystalline aluminosilicates with pores and channels of molecular dimensions [38, 39, 40, 41, 42, 43]and exhibit ion-exchange properties and absorption and reactions of molecules within their cages. They have been used for improving the performance of existing sensors or as the sensing medium itself due to their absorption, diffusion and catalytic properties, which make them excellent physical and chemical filters for enhancing selectivity [44]. The adsorption properties of zeolites have been exploited in conducting polymer-zeolite composites [45, 46, 47, 48]. For example, polymer/zeolites composites with improved sensor responses have been reported for gas sensors [49, 50, 45, 51]. In this context, employing zeolite as a functional element and combining it with graphene has great potential to achieve enhanced sensing functionalities as zeolite exhibits high surface area due to its internal microporosity, providing an excellent site for the absorption of the graphene molecules.

This study presents new insights into *in situ* fictionalisation of Kombucha pellicles to enhance their electrical properties and introduce targeted responsive capabilities. We performed electrical measurements on both unmodified and functionalised cellulose mats under mechanical stimuli using weight load application and pattern projection. The results showed that the functionalised Kombucha cellulose mats exhibit significantly higher sensitivity to external input compared to the control unmodified cellulose mats. This work contributes to the emerging field of bioelectronic sensing and establishes Kombucha cellulose as an exciting, sustainable scaffold material for information recognition and transmission.

## 2. Methods

### 2.1. Preparation of pure and functionalised Kombucha cellulose

For modulating the electrochemical properties of Kombucha cellulosic mats, graphene and zeolite were used for *in situ* functionalisation. For the production of the unmodified and functionalised Kombucha cellulose samples, a SCOBY culture of yeast and bacteria acquired from Freshly Fermented Ltd (Lee-on-the-Solent, PO13 9FU, UK) was inoculated in a sucrose tea infusion. For the infusion, black tea was selected as it has demonstrated higher levels of cellulose production [4]. The infusion was prepared with 5L boiled tap water, 500g of white granulated sugar (Tate & Lyle, UK), and 10 black tea bags (Taylors Yorkshire Teabags 125 g, UK) in a plastic container and left to reach room temperature (22°C). The SCOBY mat was then placed in the solution, and the container was stored in darkness and maintained at static conditions of 19°C. A lid with 8 × 0.5 mm holes was used to fully enclose the plastic container. Film formation was observed on the sixth day as thin layers above and below the native SCOBY, and the thickness increased with fermentation time. We replaced the solution every 12-14 days before the increase in cellulose production reached a plateau, following the same infusion protocol as mentioned above. Our rationale was that a rapid decrease of sucrose has been reported from the fifth to the fifteen-day, which is attributed to the yeast extracellular enzyme production [4] resulting from the kinetic behaviour that the Kombucha SCOBY exhibits, where yeasts hydrolyse sucrose for their own consumption, producing ethanol and carbon dioxide, and making available reducing sugars for bacterial metabolism [52, 11]. After 8 weeks, the cellulose mats were transferred to a 240×240 mm Petri dish each. 30 ml of newly made sucrose black tea solution was added to each Petri dish, instigating the fermentation process and allowing for the continuation of mat growth. After 7 days of inoculation, we proceeded with the experiments. The two cellulose mats were predominantly homogenous, with a high intensity of yellow and brown colour observed. The yellowness can be attributed to the Maillard reactions taking place during the fermentation and film conditioning occurring between the sugars and amino acids in addition to coloured phytochemicals [4]. The brown colour has been reported to be caused by melanoidins, which are coloured and nitrogen-containing compounds [53, 54]. The thickness of the mats was 3.3 mm and 4.2 mm and was measured with an electronic digital caliper (accuracy ± 0.2 mm, Vodlbov, China). The pH was monitored starting from 6.49 and reducing to 3.02 using a digital pH meter (accuracy ± 0.1% ± 2 digits, VWR pH110, Belgium).

One cellulose mat sample (thickness 3.3 mm) was functionalised with graphene and zeolite. For the functionalisation, 5.372g of graphene powder (HCG10000-P-1, Graphitene Energy Storage Series https://www.graphitene.com) were mixed with 1.289 g of Zeolite Y powder (ThermoFisher Scientific UK, 045862.36). The thickness of graphene powder was 1-3 layers with a lateral size 0.5 – 5 *μ* m, a surface area of 500 m2/g and an electrical conductivity *>*104 S/m. The surface area of zeolite Y was 927 m^2^*/*g. To achieve good interfacial bonding between the graphene and zeolite molecules, we used PTFE solvent (Sigma-Aldrich, 665880), and the mixture was stirred until a homogeneous solution was achieved (the total mixture weight was 12.965 g). The graphene-zeolite Y mixture was deposited on the surface of the Kombucha cellulose film using a razor blade and dispersed homogeneously until fully coated. After coating the top surface, the film was folded in such a manner that the mixture remained enclosed within the film. For the in situ fabrication of the Kombucha-graphene-zeolite hybrid living material, the method of microbial fermentation was used and therefore, the functionalised film was placed back again into the Petri dish to allow further growth. After 14 days, a new cellulose mat was formed, fully encapsulating the functionalised cellulose mat (Figure 1). The size of the two samples was 240×240 mm.

**Figure 1:**
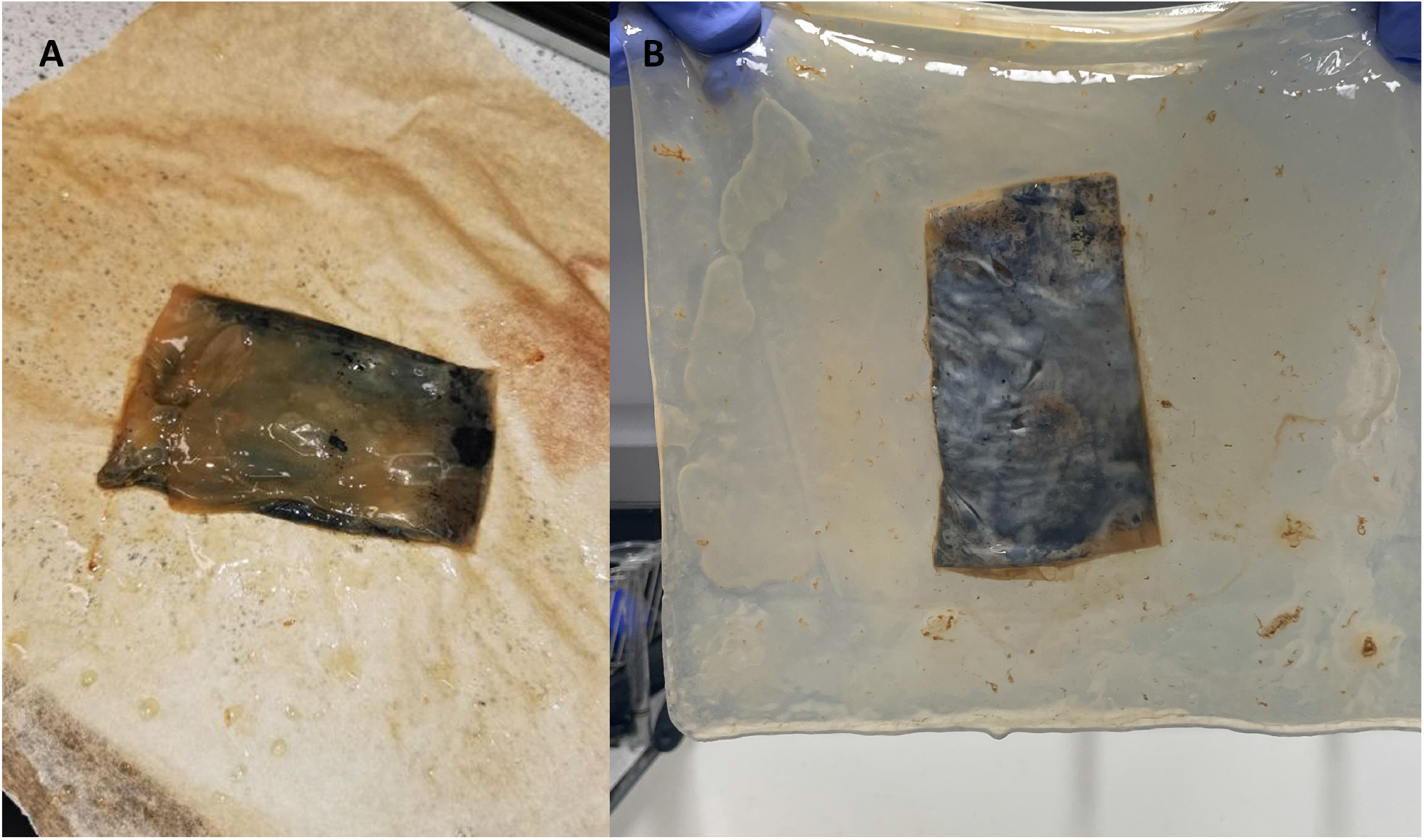
(A) The Kombucha cellulose mat after functionalisation with graphene and zeolite Y. The mat was folded to avoid spillage of the mixture when entered back to the Kombucha solution. (B) Functionalised Kombucha cellulose mat after 14 days of growth.

### 2.2. Experimental set-up and characterisation

The unmodified and functionalised Kombucha cellulose mats were placed on top of a cardboard box with dimensions of W122 × L160 mm and secured in place with plastic and metallic clamps prior to performing any measurements. To assess the response and mechanical properties of the samples when exposed to compression, we used load application. For the mechanical stimulation of the samples, manually cut square aluminium weights of 100g (W51 × L51 × H14.2 mm), 200g (W51 × L51 × H28.4 mm), 300g (W51 × L51 × H42.7 mm) and circular aluminium weights of 100g (D51 × H18.1 mm), 200g (D51 × H36 mm), 300g (D51 × H54.4 mm) were employed. The weights were applied to both samples in different combinations to attain loads of 100g, 200g, 300g, 400g and 600g (Figure 2). Both patterns of weights (square and circular) were applied to the two cellulose samples.

**Figure 2:**
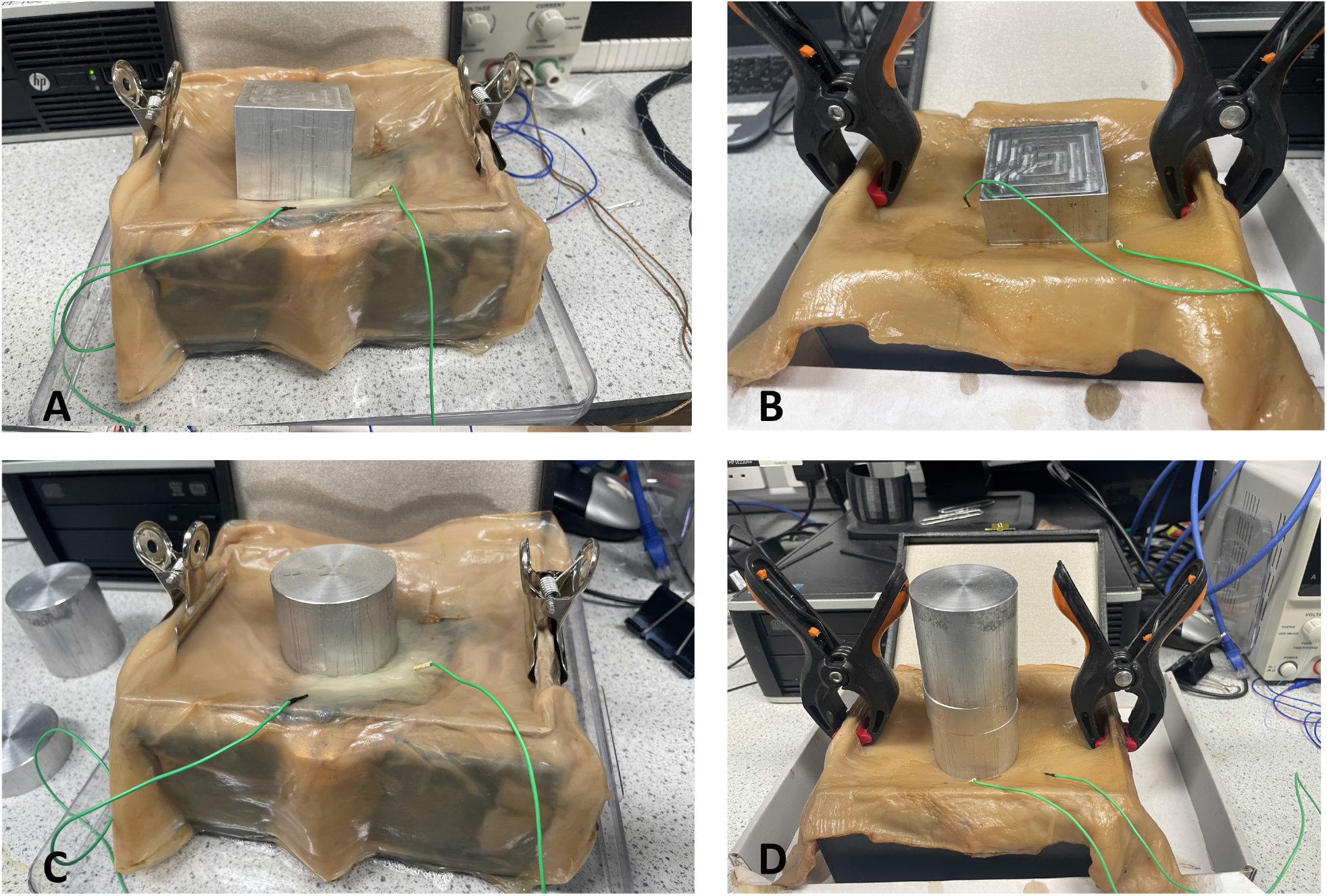
(A), (C) Impedance, capacitance, resistance and I-V measurements taken from functionalised Kombucha cellulose mat, 240 × 240 mm size, under square and circular weights of 100 g, 200 g, 300 g and 600 g. (B), (D) Impedance, capacitance, resistance and I-V measurements taken from functionalised Kombucha cellulose mat, 240×240 mm size, under square and circular weights of 100 g, 200 g, 300 g and 600 g.

Electrical characterisation of the unmodified and functionalised mats prior to and during the application of 100g, 200g, 300g and 600g loads were performed. A pair of iridium-coated stainless steel sub-dermal needle electrodes (Spes Medica S.r.l., Italy), with twisted cables, was pierced through the mats with a distance between the electrodes of 50-60mm. A Keithley 2450 SourceMeter was used for the provision of high-precision current-voltage (I–V) sweep measurements (type linear dual, voltage range from −1V to 1V, step 10mV, count finite 20, source limit 1A). Impedance, capacitance and resistance measurements were taken using a BK Precision LCR Meter (model 891) within the frequency range of 20Hz and 300kHz. The experiments were repeated to ensure error elimination.

For the recordings of the electrical activity before and after the weight application, an ADC-24 (Pico Technology, UK) high-resolution data logger with a 24-bit A/D converter was used. Weights of square and circular patterns of 100g, 300g and 400 g were applied to the samples. 8 pairs of iridium-coated stainless steel sub-dermal needle electrodes were inserted in the samples; the first 4 (Channels 1-2, 3-4, 5-6, 7-8) in the unmodified cellulose sample and the remaining 4 (Channels 9-10, 11-12, 13-14, 15-16) in the functionalised sample (Figure 3). We recorded electrical activity simultaneously, one sample per second and each pair of electrodes reported a difference in the electrical potential between the electrodes. All readings were taken at a room temperature of 19°C. During the recording, the logger undertook as many measurements as possible (typically up to 600 per second) and saved the average value. The distance between electrodes was 10-20 mm.

**Figure 3:**
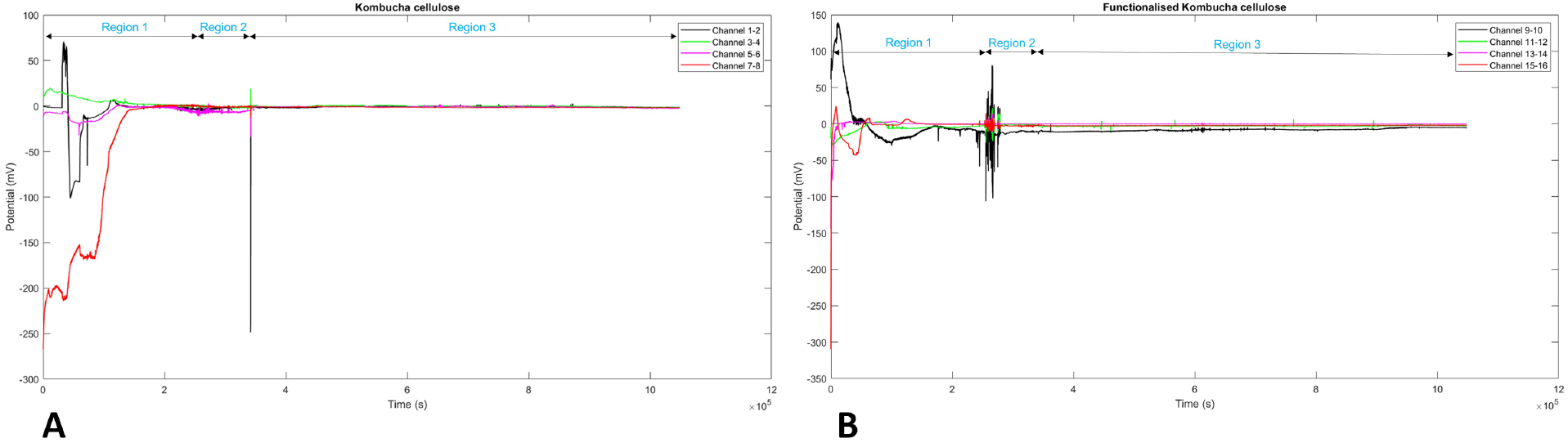
Spiking activity of 240 × 240 mm samples when exposed to different loads. The graphs display the electrical activity over time, measured in potential (mV) versus time (sec), from both Kombucha (Ch1-Ch4) and Kombucha with graphene-zeolite Y cellulose mats (Ch5 –Ch8). In (A), Region 1 shows the application of a 100 g box load, Region 2 represents the application of a 400 g box load, and Region 3 represents the application of a 300 g cylinder load. In (B), Region 1 represents the application of a 100 g cylinder load, Region 2 represents the application of a 400 g cylinder load, and Region 3 represents the application of a 300 g box load.

The microstructure of the unmodified and functionalised Kombucha cellulose samples was analysed by scanning electron microscopy and environmental scanning electron microscopy (FEI Quanta 650). For the preparation prior to imaging, the samples were air-dried and coated with a gold layer using an Emscope SC500 gold sputter coating unit. Images were acquired with a magnification range of 5000x - 20000x and a high voltage of 2 kV. To the best of our knowledge, this is the first time that Kombucha cellulose has been functionalised to evaluate its electrical properties and response to load application.

## 3. Results

### 3.1. Morphological and chemical characterisation

SEM imaging performed on pure and functionalised Kombucha-graphene-zeolite samples demonstrated that the structure of the pure Kombucha mat changes significantly with the integration of the graphene-zeolite particles. The morphological features of pure Kombucha reveal that the microbial community of yeast and bacterial cells present mostly ellipsoidal structures, are compactly and uniformly dispersed, and are entangled into a dense mesh of cellulosic fibres that form three-dimensional web-like structures. The presence of multiple-size yeast cells is observed, spanning from 1.93*μ*m to 7.76*μ*m in length. Chain yeast elongations are visible, and protrusions on the cell surfaces show cell budding mechanisms [55]. Figure (Figure 4) shows bud and birth scars as well as bud-parent junctions (marked with red circles, yellow and red arrows accordingly). Multipolar budding is also clearly identified on some cells (marked with a yellow circle).

**Figure 4:**
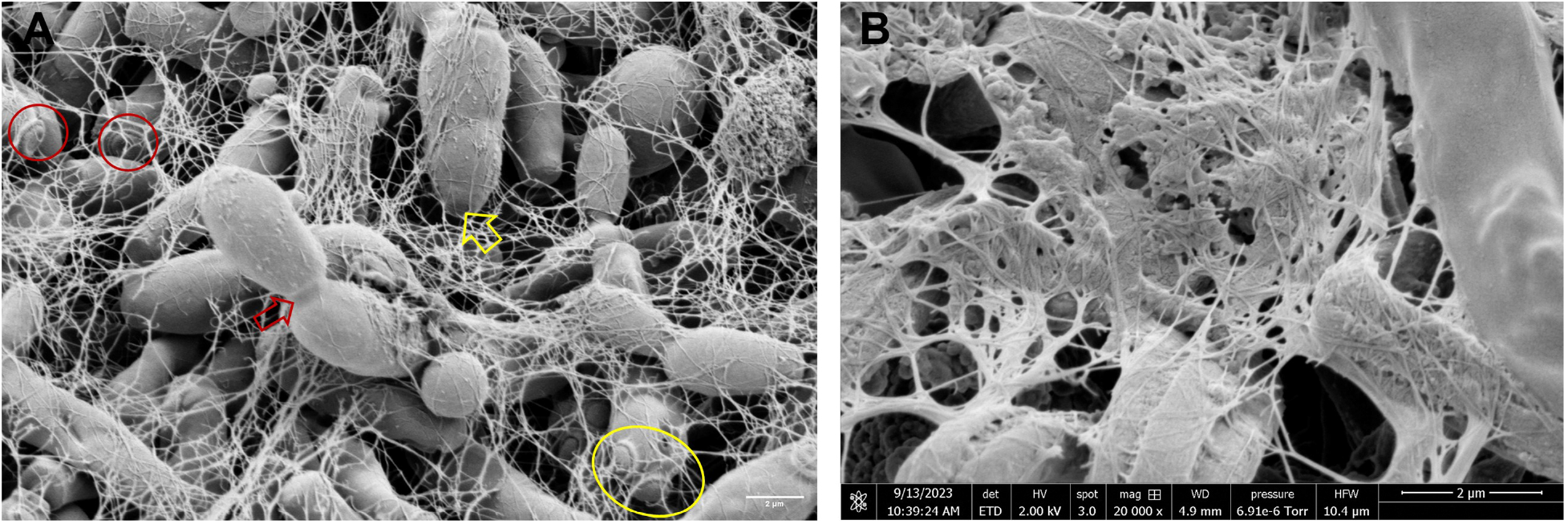
SEM images of pure Kombucha mat. A. The presence of multiple-size yeast cells is identified, with lengths ranging from 1.93*μ*m to 7.76*μ*m. Cell bud scars, birth scars, multipolar budding and bud-parent sites are marked in the image with red circles, yellow and red arrows and a yellow circle accordingly. Image magnification is 8000x. B.A network of cellulosic fibres with a branching structure. Image magnification is 20000x.

Images of the functionalised sample revealed dense aggregations of zeolite crystals with morphologies of spherical agglomerates of tiny crystals next to larger polyhedrons surrounded by typical flakes of graphene particles [56] and integrated with scattered Kombucha microbes. The presented high-density configurations may be attributed to the high microporosity of zeolite active sites, which provides space for the adsorption and exchange of cations, leading to the attachment of the graphene molecules. This connection could favour electron transport and contribute to the electrochemical performance of the functionalised Kombucha film. In addition to this, the cellulose fiber network appears to be less dense and finer where the graphene-zeolite formations are present. The measured cluster assemblies have a diameter of 9.61*μ*m and 7.75*μ*m (see Figure 5). Contrary to pure Kombucha images, microbial cells appear in lower densities in the functionalised Kombucha images, but elongations and birth scars are still present, suggesting that growth and replication mechanisms continue to take place after functionalisation. The above findings indicate that the new structural formations may involve the microscopic movement of the microbial population due to the mechanical and physical forces built up from the graphene-zeolite encapsulation during in situ functionalisation. Moreover, it is observed that the graphene-zeolite particles are found entangled and attached with the microbial cells and in parts enveloped by the cells, suggesting integration of the living biological and synthetic components and, therefore, the possible synthetic morphogenesis of a new and higher-ordered structure. This may open up remarkable opportunities for the in-situ development of functionalised hybrid materials that can self-assemble, self-replicate, self-generate and self-grow after the fusion of their living and synthetic components. Energy Dispersive Spectroscopy (EDS) shows the distribution of elements within the functionalised Kombucha film(Figure 6). In the material distribution analysis, very low Al content is observed, demonstrating high hydrothermal stability [57].

**Figure 5:**
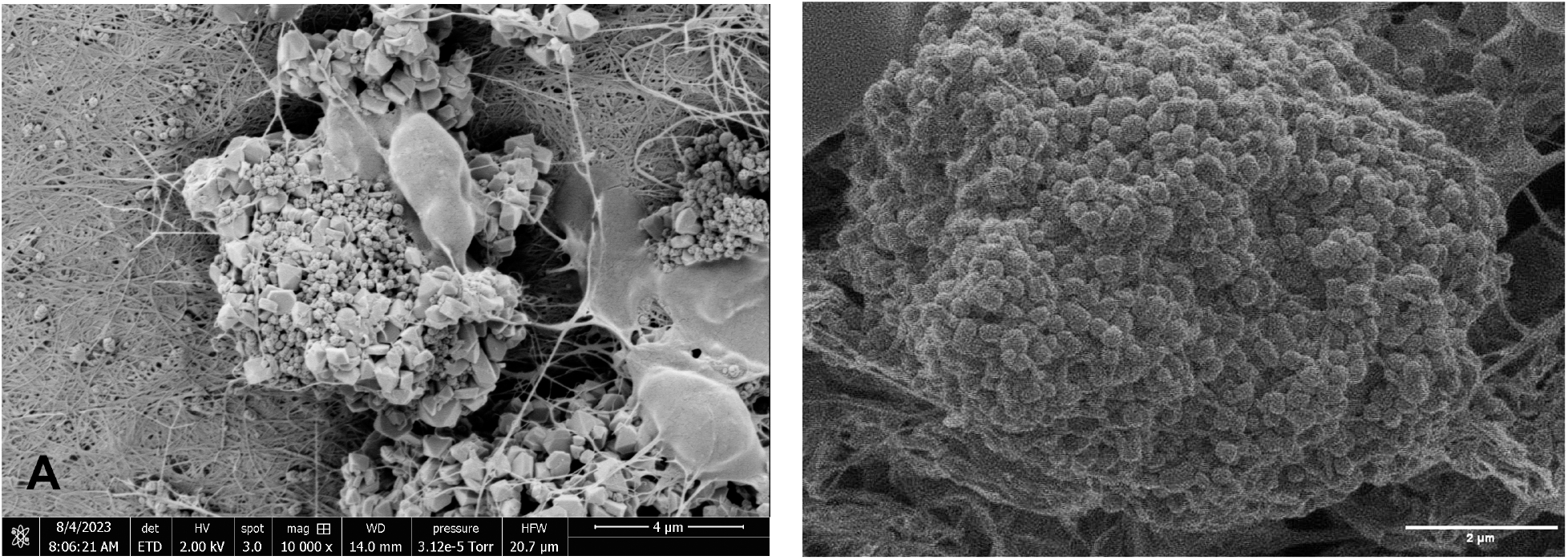
SEM images of functionalised Kombucha mat. A. Clusters of zeolite crystals and graphene particles entangled with microbes. Image magnification is 10000x. B. Higher magnification image of graphene-zeolite assembly formations discern graphene flakes attached to zeolite crystals. Image magnification is 20000x.

**Figure 6:**
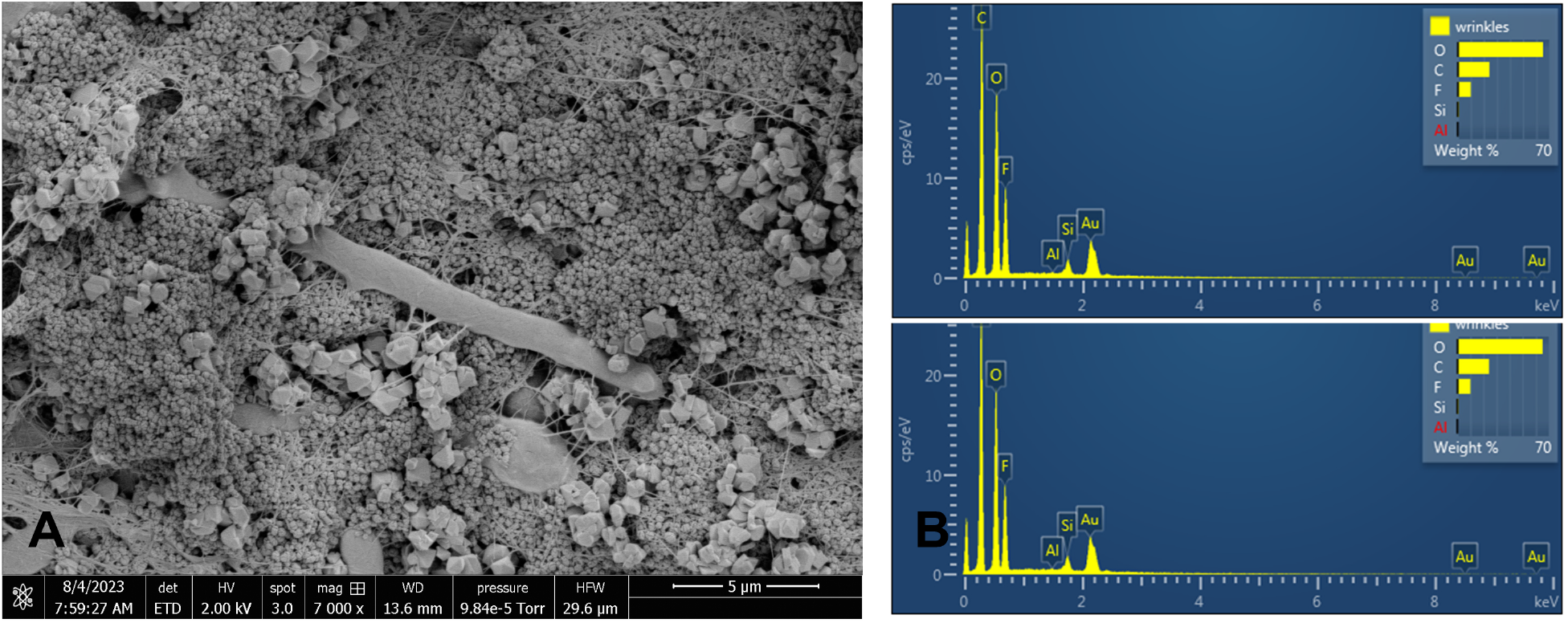
(A) Yeast cells integrated into zeolite-graphene nanoparticles. Image magnification is 7000x. (B) EDS graphs showing the distribution of elements for the functionalised Kombucha sample.

### 3.2. DC and impedance properties

Measurements have been carried out in the frequency range between 1 and 300 kHz. Series resistance as a function of the frequency is shown in Figure 7. The upper row shows the Kombucha mat in different conditions (dry, wet and unloaded, and with cylinder and box loads from 100 to 600 grams), while the bottom row shows functionalised mats. The dry condition is such that the functionalised mat features a resistance one order of magnitude lower than the pristine Kombucha mat due to the enhancement provided by graphene addition. The consecutive loading of the mat produces a reduction of the resistance that achieves a maximum with the intermediate loading of 200 g and then recovers back and eventually becomes even higher than the unloaded measure for the loading of 600 g. This effect might be due to pseudoelastic stretching of the matrix under the normal forces exerted by the loads, with concurrent reduction of the section available for conduction. A remarkably similar trend might be found in the functionalised Kombucha mat, where the 100 and 200 g loads reduce the resistance, and the 300 and 600 ones increase it, yet to values lower than the unloaded case. About the shape of the load, we can notice that the cylinder weight can produce very pronounced responses, particularly in the load range of 200 - 300 g, for both mats.

**Figure 7:**
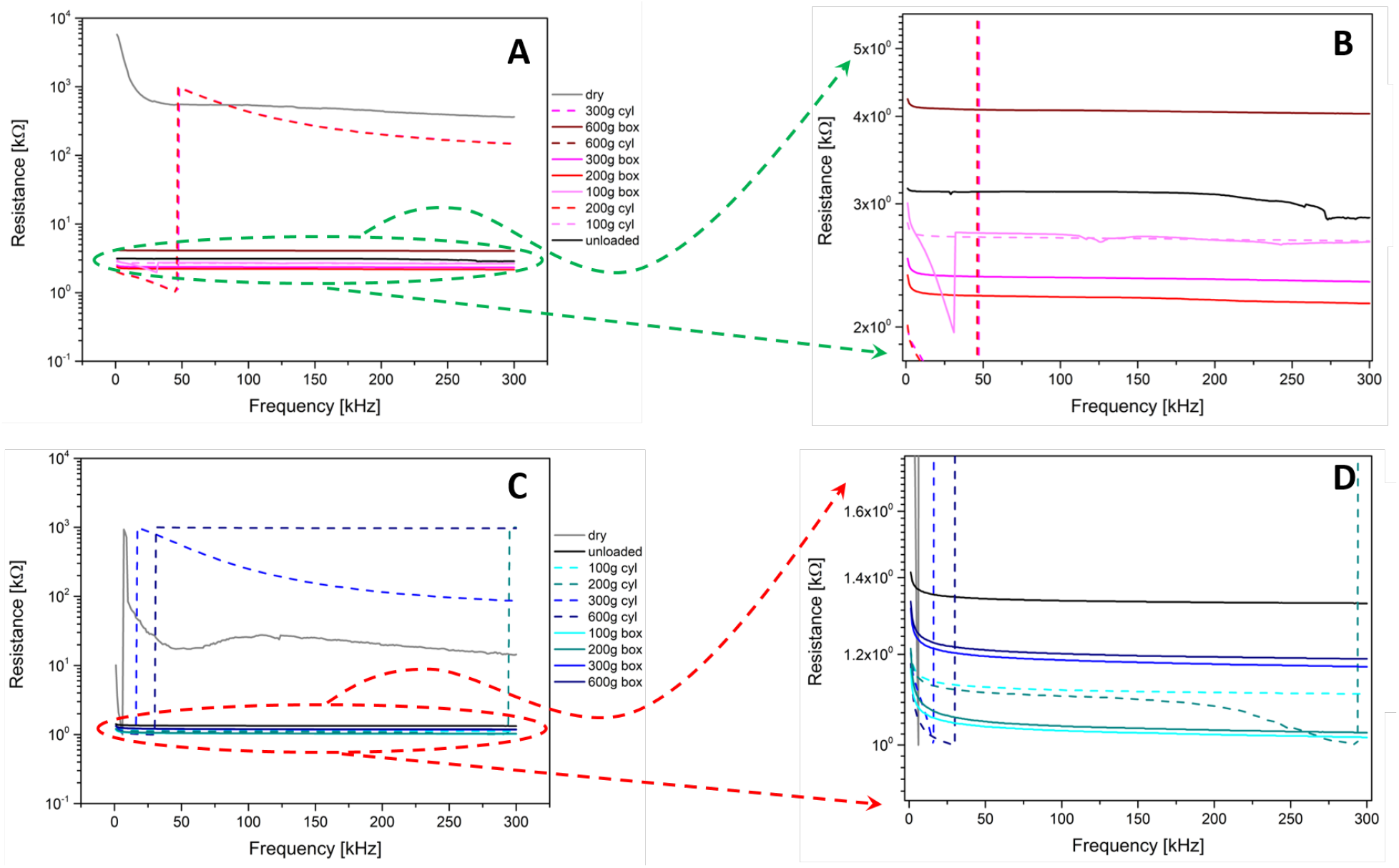
Series resistance as a function of the frequency. (A) Kombucha response to the different loads and (B) inset showing a zoom of the circled area. (C) The functionalised Kombucha response to the different loads and (D) inset showing a zoom of the circled area.

In Figure 8, the capacitance is shown in the same frequency range discussed above. The unloaded curve of the pristine mat versus the functionalised one shows that normally, the capacitance of the latter is approximately double that of the former, which can be due to the addition of zeolite compounds having a higher dielectric constant. Nevertheless, we should remember how much water content in a mat can influence this aspect, as the dielectric constant of water is very high: the dry mat curve shows how small its capacitance can be in the absence of water. Looking at the pristine mat curves, we cannot infer any specific pattern: the measurements cannot correlate with the load amount and/or its shape. On the contrary, the functionalised mat curves feature a more controlled behaviour that perfectly maps the observations based on the resistance previously discussed: the deformation produced by 100 and 200 g loads reduces the capacitance, as the displacement of the hydrogel under the force exerted by the load reduces the surface of the dielectric layer incorporated in the capacitor. Other phenomena might occur, such as the capacitive coupling with the load materials (i.e., aluminium and polymers), making it difficult to infer the shape of the load from the measurements.

**Figure 8:**
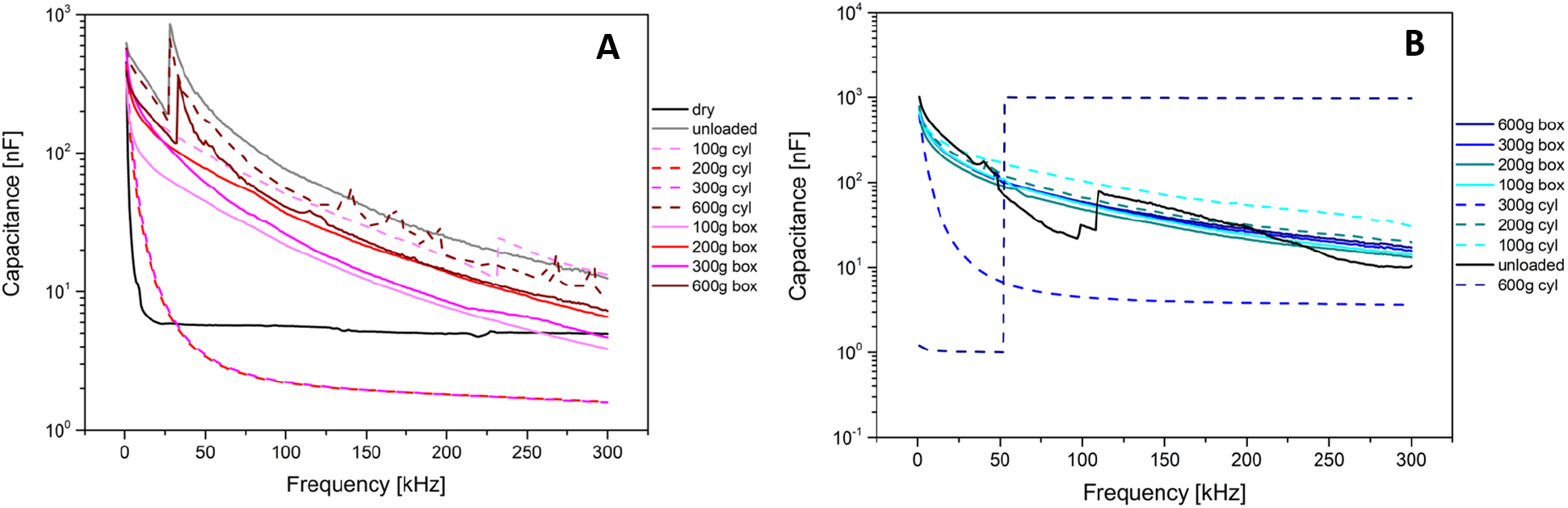
Capacitance as a function of the frequency. (A) Kombucha response to the different loads and (B) functionalised Kombucha response to the different loads.

Another important analysis performed is the collection of IV curves along multiple hysteresis cycles to put in evidence typical current ranges and eventually a memristive behaviour. Figure 9 shows that in the 1 V range, the currents are of the order of 10 *μ*V for both the pristine and functionalised mats, but in the former case, all curves are very close, and the loading does not provide any strong variation, while in the latter we can see how much the loaded curves differ from the unloaded one, having currents five times bigger. Therefore, loading the functionalised mat produces higher currents (lower resistances), while loading the pristine mat produces lower currents (higher resistances). An enhanced conductivity can explain this effect due to the percolation of the fillers added to the functionalised mat. The higher pressure creates more contact points between inorganic conductive fillers, such as the graphene flakes, and reduces the mat resistance.

**Figure 9:**
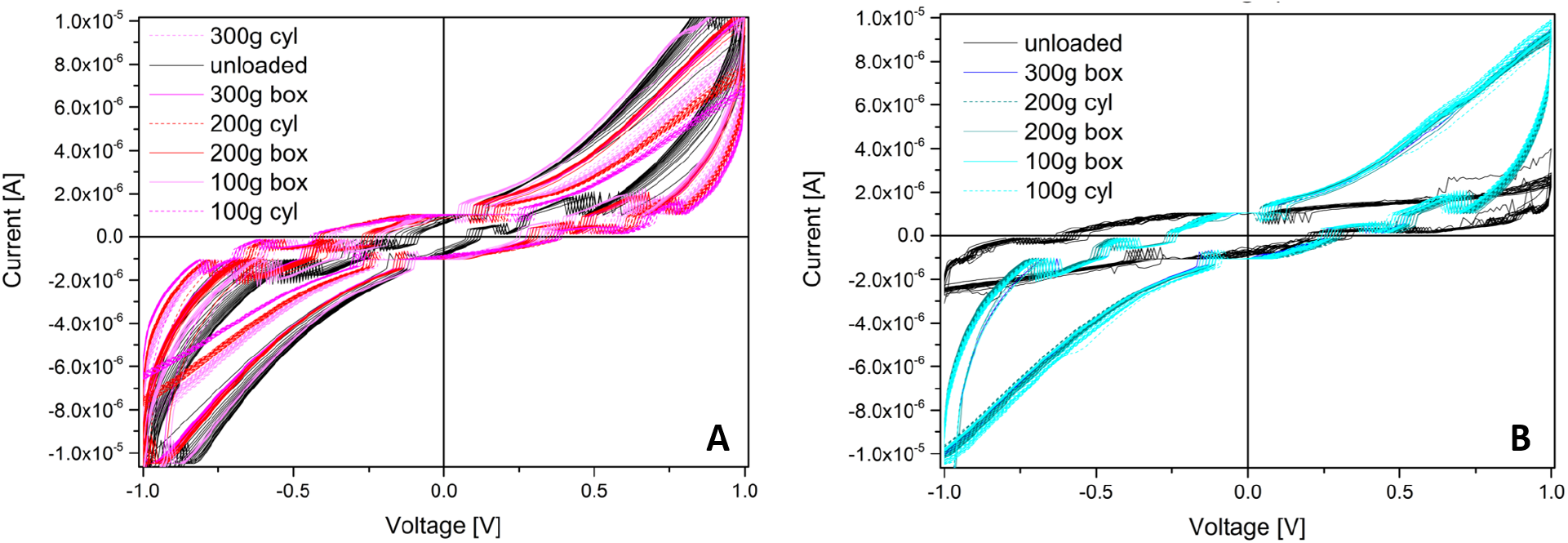
Hysteresis cycles showing the IV response of the mats. (A) Kombucha response to the different loads and (B) functionalised Kombucha response to the different loads.

### 3.3. Spiking activity and higher order complexity analyses

The spontaneous spiking activity of two mats was continuously recorded while loads were being applied. The recorded profiles from four differential channels are shown in Figure 10 for the entire measurement period of 11 days. By examining each channel, we can observe a certain degree of correlation involving transitions and increased spiking activity with remarkably similar characteristics. Typically, the spontaneous activity range is around 20 mV. However, some channels exhibit a large baseline fluctuation of about 175 mV (for example, channels 1-2 and 5-6 of the pristine mat) and approximately 200 mV (for example, channels 9-10 of the functionalised mat). To gain a clearer understanding of what is happening in various situations, we used the Fast Fourier Transform (FFT) and Higher Order Complexity (HOC) metrics [58, 59] as shown in Figure 11.

**Figure 10:**
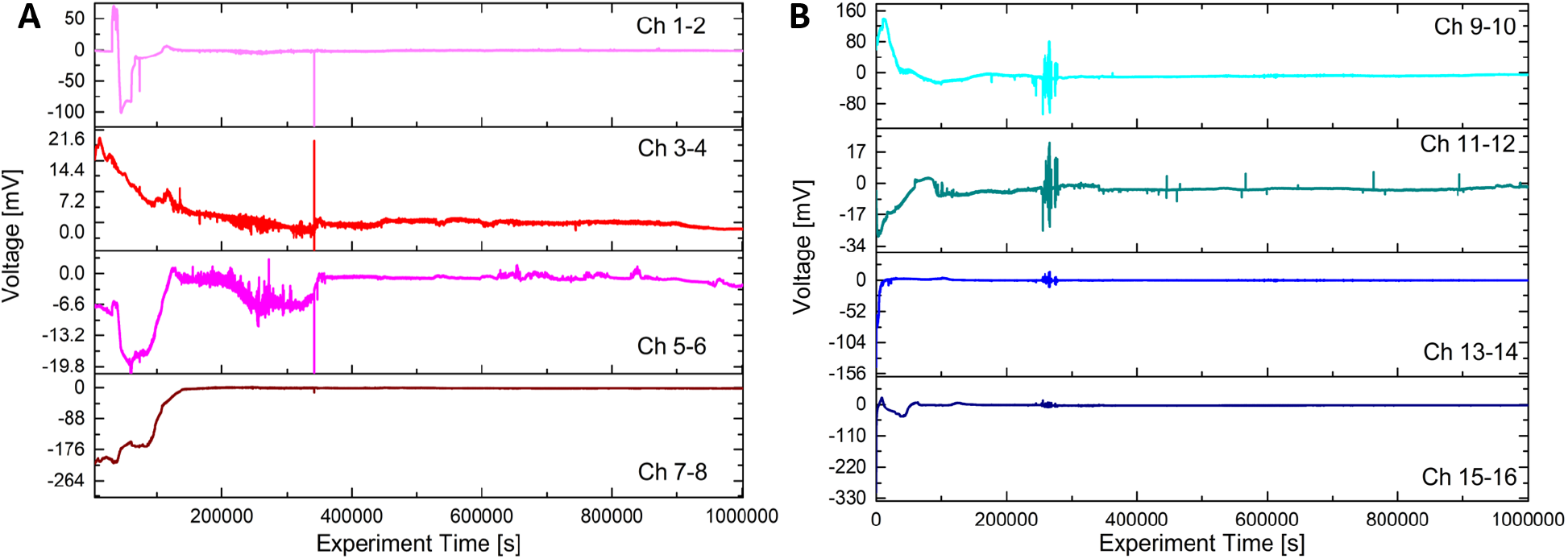
Recordings showing the spiking response of the mats. (A) Kombucha response to the different loads and (B) Functionalised Kombucha response to the different loads.

**Figure 11:**
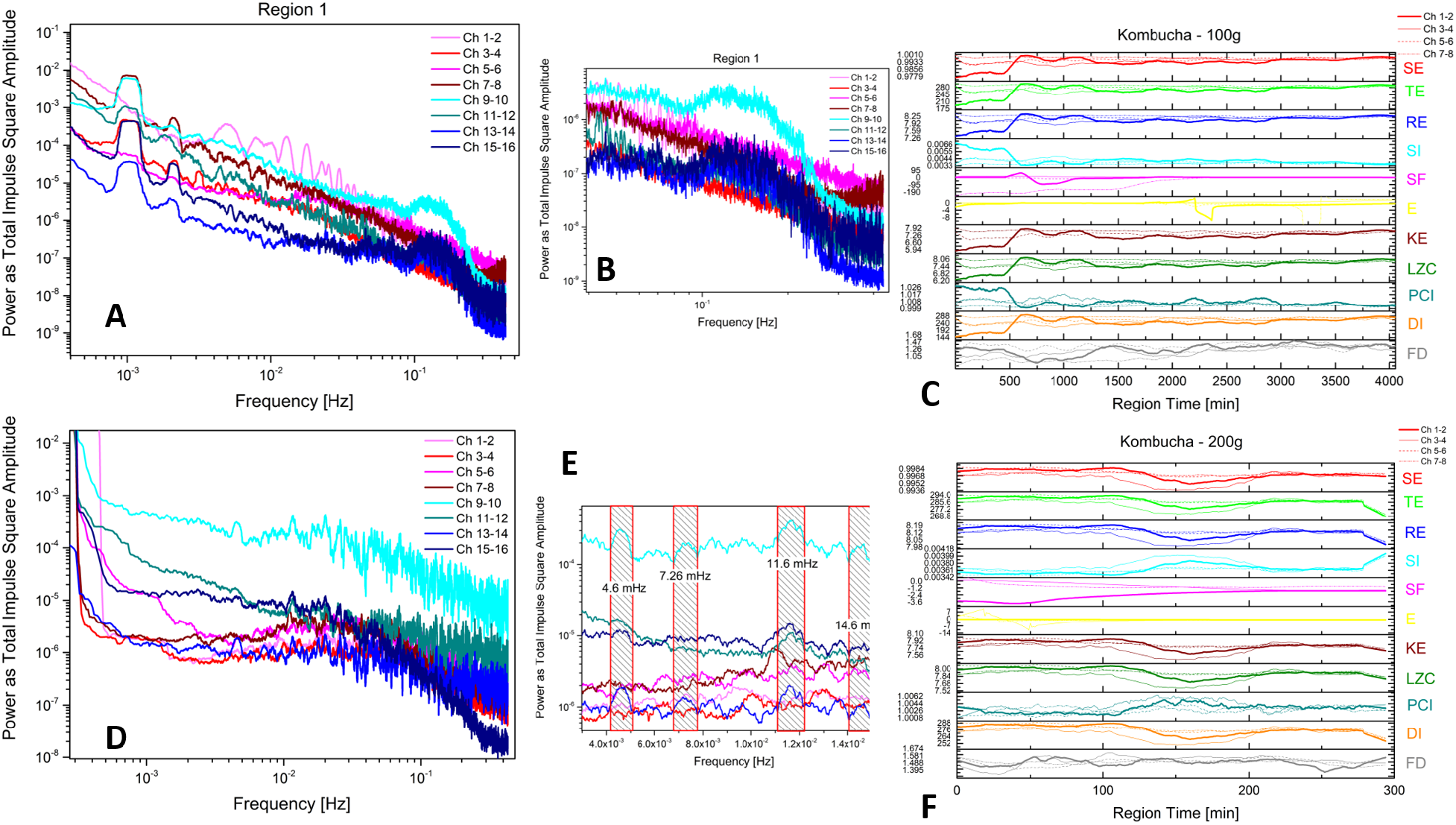
Regional details of the spiking activity features. (A) The Fast Fourier Transforms correspond to 100 g loading of the Kombucha mat and functionalised Kombucha mat. (B) A zoomed-in view highlights a broad peak structure. (C) Higher-order complexity measures for the same region have been provided, pristine mat. SE stands for Shannon entropy, TE represents Tsallis entropy, RE stands for Rényi entropy, SI represents Simpson Index, SF represents Space Filling, E represents Expressiveness, KE represents Kolmogorov complexity, LZC represents Lempel-Ziv complexity, PCI represents Permuted Complexity Index, DI represents Diversity Index, and FD represents Fractal Dimension. (D) The Fast Fourier Transforms for the region corresponding to 200 g loading of the Kombucha mat and functionalised Kombucha have been shown. (E) A zoomed-in view highlights a broad peak structure. (F) Higher-order complexity measures for the same region, pristine mat, have been provided.

The FFT analysis shows how the noise spectrum across the mat sensors is distributed over different frequencies, ranging from approximately 1 mHz up to 0.5 Hz, corresponding to half the sampling frequency. The first row of graphs depicts data from “Region 1” when a 100 g load was positioned on the mats. The red lines (differential electrode couples 1-2, 3-4, 5-6 and 7-8) represent the pristine Kombucha, while the blue lines (differential electrode couples 9-10, 11-12, 13-14 and 15-16) represent the functionalised Kombucha. A fundamental oscillation mode occurs across all channels at 1 mHz, with visible superior harmonics at 2, 3, and 4 mHz. It is unclear whether the source of this ultra-low frequency mode is an external noise disturbance or part of the mat’s electrical oscillations. Notably, there is a broad peak spanning 0.1 to 0.2 Hz, a genuine signal component from the functionalised mat not observable in the pristine mat data. The complexity measures display consistent behaviour where the curves either show a lower plateau for the first 500 minutes followed by recovery to higher levels for the remaining 4000 minutes (SE, TE, RE, SF, KE, LZC, DI) or an initial higher plateau followed by decay to lower values (SI, PCI, FD). This demonstrates that implementing proper higher-order complexity (HOC) measures makes it feasible to interpret the mat’s spontaneous oscillations as a sensor response with long timescales.

Examining data from “Region 2,” where a 200 g load was applied for 300 minutes, the shorter duration limits comparability to the complexity trends in Region 1. However, noteworthy FFT responses occur, including cleaner oscillation profiles and specific modes at 4.6, 7.3, 11.6 and 14.6 mHz visible on the pristine mat. Similar modes at slightly shifted frequencies also emerge on the functionalised mat, generally skewed towards faster processes, potentially indicative of increased conductivity.

Further analysis using Short Time Fourier Transforms (STFT) generates maps with colour-coded intensity fluctuations over time and frequency, depicted in Figure 12. The top row shows the STFT for a functionalised mat differential couple (Region 1). A specific 5-hour period exhibits multiple harmonics and a complex structure circled with a dashed ellipse and magnified to the right, resembling a vocal signal with fundamentals around 40 mHz and harmonics approaching 200 mHz. Interestingly, a comparable 1-hour fragment occurs in Region 2 data from the pristine mat with fundamentals around 30 mHz and harmonics near 250 mHz. Audio files were generated by the sonification of these recordings along with another excerpt from Region 3 using Melobytes software (Gardos Software Ltd. https://melobytes.com). The mp3 files are provided as supporting materials.

**Figure 12:**
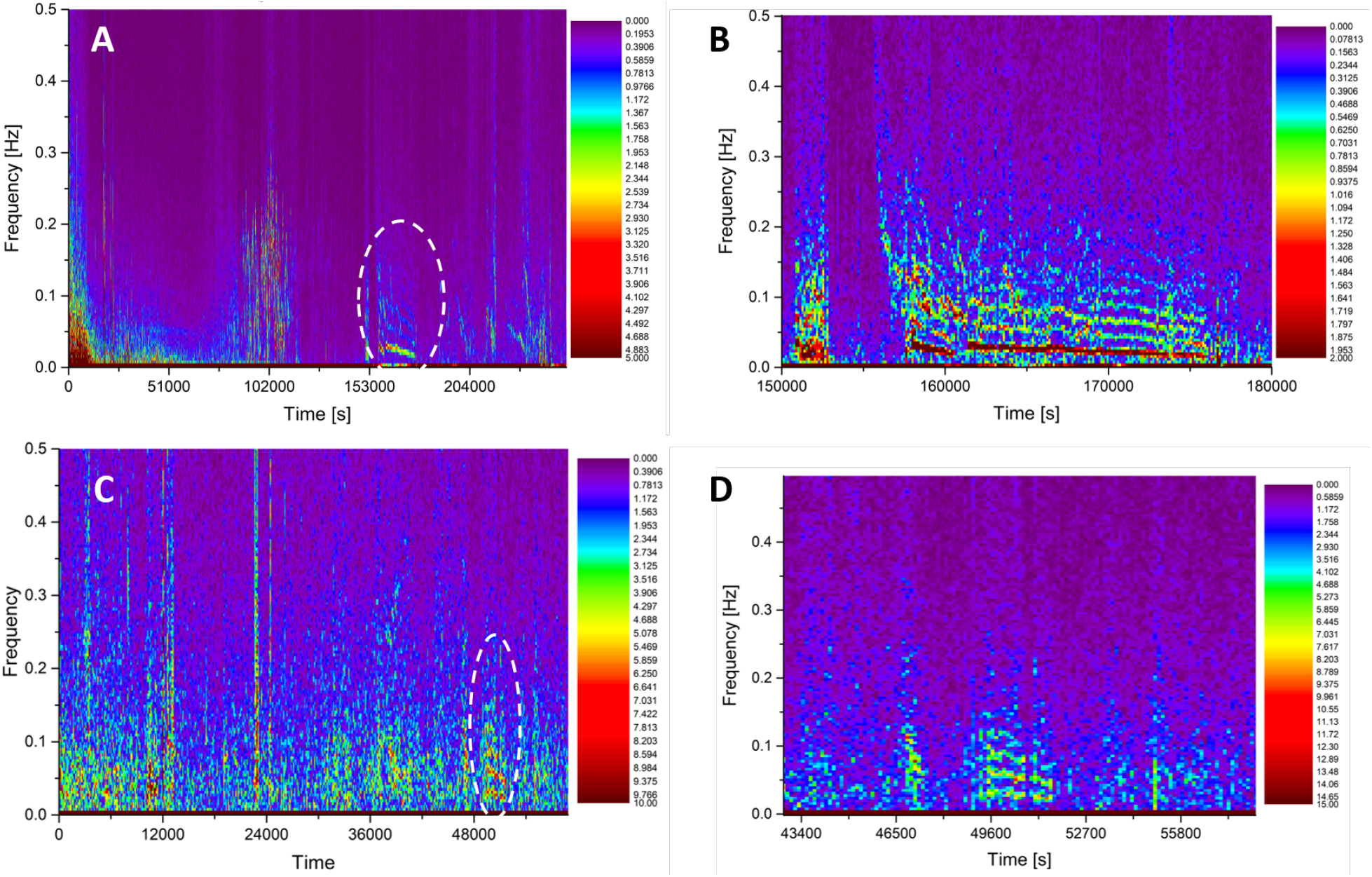
Regional details of the spiking activity features. (A) Short-time Fast Fourier Transforms on the region corresponding to 100 g loading of the differential channel 13-14 of the functionalized Kombucha mat. (B) A zoomed-in view highlights a peculiar feature, as indicated by the ellipse. (C) Short-time Fast Fourier Transforms on the region corresponding to 100 g loading of the differential channel 7-8 of the pristine Kombucha mat. (D) A zoomed-in view highlights a peculiar feature, as indicated by the ellipse.

## 4. Conclusion

In this study, we have demonstrated that Kombucha zoogleal mats, functionalised with graphene and zeolite particles, exhibit unique electrical properties. Two key findings are memfractive behaviour and distinctive electrical spiking in response to mechanical loading.

Memfractive materials combine properties of memristors, memcapacitors, and meminductors [60]. As a memristive material implied by its material nature [61, 62, 63, 64], Kombucha mats could enable various logic circuits [65], stateful logic operations [66], memory-aided logic circuits [67], logic operations within passive crossbar arrays of memristors [68], memory-aided logic circuits [67], self-programmable logic circuits [69], and memory devices [70]. Their memfractive nature also makes the mats suitable for integration into diverse memory and computing devices, including biocompatible electronics and bio-wearables.

The observed electrical spiking from mechanical stimulation (i.e., apply loads) could be harnessed to create biocompatible sensors. Incorporating the mats into wearable devices or implantable sensors may facilitate real-time detection of physiological changes in the human body or other biological systems with potential applications in real-time health monitoring. The electrical spiking could also be explored for energy harvesting purposes. If the spiking can be converted into a usable form of energy, it might contribute to powering small electronic devices or sensors in remote or hard-to-reach locations. Combining Kombucha zoogleal mats with functional nanoparticles could lead to hybrid systems that leverage the strengths of both biological and synthetic elements.

This work demonstrates the early-stage potential for hybrid systems combining Kombucha zoogleal mats with functional nanoparticles to leverage both biological and synthetic elements. This interdisciplinary approach could open up new possibilities in sustainable electronics and computing applications that are impossible with either system alone. Specifically, the biodegradable nature of Kombucha materials makes them well-suited for environmentally friendly devices and components aimed at reducing e-waste. Devices or components that naturally decompose after use could reduce electronic waste and positively impact sustainability.

## Competing interests

All authors declare no competing interests.

## Acknowledgements

Authors are grateful to David Patton for assisting with SEM imaging and Dr Jun Yao for her advice on graphene and zeolites.

## Funding

This research did not receive any specific grant from funding agencies in the public, commercial, or not-for-profit sectors.

## Data availability

The data is accessible via the online database Zenondo and can be accessed via the link https://doi.org/10.5281/zenodo.10445665).

## Notes

### Competing Interest Statement

The authors have declared no competing interest.

## References

[1] M. Iguchi, S. Yamanaka, A. Budhiono, Bacterial cellulose—a masterpiece of nature’s arts, Journal of materials science 35 (2) (2000) 261–270.

[2] M. M. Abeer, M. C. I. Mohd Amin, C. Martin, A review of bacterial cellulose-based drug delivery systems: their biochemistry, current approaches and future prospects, Journal of Pharmacy and Pharmacology 66 (8) (2014) 1047–1061.

[3] B. Thomas, M. C. Raj, J. Joy, A. Moores, G. L. Drisko, C. Sanchez, Nanocellulose, a versatile green platform: from biosources to materials and their applications, Chemical reviews 118 (24) (2018) 11575–11625.

[4] Y. A. R. Tapias, M. V. Di Monte, M. A. Peltzer, A. G. Salvay, Bacterial cellulose films production by kombucha symbiotic community cultured on different herbal infusions, Food Chemistry 372 (2022) 131346.

[5] Y. Yamada, P. Yukphan, Genera and species in acetic acid bacteria, International journal of food microbiology 125 (1) (2008) 15–24.

[6] A. Singh, K. T. Walker, R. Ledesma-Amaro, T. Ellis, Engineering bacterial cellulose by synthetic biology, International Journal of Molecular Sciences 21 (23) (2020) 9185.

[7] C. Cottet, Y. A. Ramirez-Tapias, J. F. Delgado, O. de la Osa, A. G. Salvay, M. A. Peltzer, Biobased materials from microbial biomass and its derivatives, Materials 13 (6) (2020) 1263.

[8] D. Laavanya, S. Shirkole, P. Balasubramanian, Current challenges, applications and future perspectives of scoby cellulose of kombucha fermentation, Journal of Cleaner Production 295 (2021) 126454.

[9] A. Chiolerio, A. Adamatzky, Acetobacter biofilm: Electronic characterization and reactive transduction of pressure, ACS Biomaterials Science & Engineering 7 (4) (2021) 1651–1662.

[10] A. Chiolerio, M. M. Dehshibi, D. Manfredi, A. Adamatzky, Living wearables: Bacterial reactive glove, Biosystems (2022) 104691.

[11] A. May, S. Narayanan, J. Alcock, A. Varsani, C. Maley, A. Aktipis, Kombucha: a novel model system for cooperation and conflict in a complex multi-species microbial ecosystem, PeerJ 7 (2019) e7565.

[12] R. M. D. Coelho, A. L. de Almeida, R. Q. G. do Amaral, R. N. da Mota, P. H. M. de Sousa Kombucha, International Journal of Gastronomy and Food Science 22 (2020) 100272.

[13] A. B. Delatorre, T. R. L. Ribeiro, N. M. C. Arcênio, G. P. B. Mothé, Estudo sobre a adequação da normativa 41 para a produção e comercialização de kombucha: estudo de caso em uma empresa de macaé, in: Congresso de Ensino Pesquisa e Extensão-CONEPE, 2020.

[14] S. Torgbo, P. Sukyai, Bacterial cellulose-based scaffold materials for bone tissue engineering, Applied Materials Today 11 (2018) 34–49.

[15] S. Patra, M. Singh, D. Pareek, K. Wasnik, P. S. Gupta, P. Paik, Advances in the development of biodegradable polymeric materials for biomedical applications (2022).

[16] A. Adamatzky, Electrical potential spiking of kombucha zoogleal mats: A symbiotic community of bacteria and yeasts, Bioelectricity 5 (2) (2023) 99–108.

[17] A. Adamatzky, G. Tarabella, N. Phillips, A. Chiolerio, P. D’Angelo, A. Nikolaidou, G. C. Sirakoulis, Kombucha electronics: electronic circuits on kombucha mats, Scientific Reports 13 (9367) (2023).

[18] H. Bai, G. Shi, Gas sensors based on conducting polymers, Sensors 7 (3) (2007) 267–307.

[19] B. Adhikari, S. Majumdar, Polymers in sensor applications, Progress in polymer science 29 (7) (2004) 699–766.

[20] T. Ahuja, D. Kumar, et al., Recent progress in the development of nano-structured conducting polymers/nanocomposites for sensor applications, Sensors and Actuators B: Chemical 136 (1) (2009) 275–286.

[21] K. S. Novoselov, A. K. Geim, S. V. Morozov, D.-e. Jiang, Y. Zhang, S. V. Dubonos, I. V. Grigorieva, A. A. Firsov, Electric field effect in atomically thin carbon films, science 306 (5696) (2004) 666–669.

[22] D. Dreyer, S.; park, cw; r, bielawski.; and ruoff, rs “the chemistry of graphene oxide, Chemical Society reviews 39 (1) (2010) 228–240.

[23] G. Wang, J. Yang, J. Park, X. Gou, B. Wang, H. Liu, J. Yao, Facile synthesis and characterization of graphene nanosheets, The Journal of Physical Chemistry C 112 (22) (2008) 8192–8195.

[24] X. Li, X. Wang, L. Zhang, S. Lee, H. Dai, Chemically derived, ultrasmooth graphene nanoribbon semiconductors, science 319 (5867) (2008) 1229–1232.

[25] P. Blake, P. D. Brimicombe, R. R. Nair, T. J. Booth, D. Jiang, F. Schedin, L. A. Ponomarenko, S. V. Morozov, H. F. Gleeson, E. W. Hill, et al., Graphene-based liquid crystal device, Nano letters 8 (6) (2008) 1704–1708.

[26] S. Ansari, E. P. Giannelis, Functionalized graphene sheet—poly (vinylidene fluoride) conductive nanocomposites, Journal of Polymer Science Part B: Polymer Physics 47 (9) (2009) 888–897.

[27] T. Ramanathan, A. Abdala, S. Stankovich, D. Dikin, M. Herrera-Alonso, R. Piner, D. Adamson, H. Schniepp, X. Chen, R. Ruoff, et al., Functionalized graphene sheets for polymer nanocomposites, Nature nanotechnology 3 (6) (2008) 327–331.

[28] S. Stankovich, D. A. Dikin, G. H. Dommett, K. M. Kohlhaas, E. J. Zimney, E. A. Stach, R. D. Piner, S. T. Nguyen, R. S. Ruoff, Graphene-based composite materials, nature 442 (7100) (2006) 282–286.

[29] H. Fan, L. Wang, K. Zhao, N. Li, Z. Shi, Z. Ge, Z. Jin, Fabrication, mechanical properties, and biocompatibility of graphene-reinforced chitosan composites, Biomacromolecules 11 (9) (2010) 2345–2351.

[30] K. Zhang, L. L. Zhang, X. Zhao, J. Wu, Graphene/polyaniline nanofiber composites as supercapacitor electrodes, Chemistry of Materials 22 (4) (2010) 1392–1401.

[31] X. Zhao, Q. Zhang, D. Chen, P. Lu, Enhanced mechanical properties of graphene-based poly (vinyl alcohol) composites, Macromolecules 43 (5) (2010) 2357–2363.

[32] T. Kuila, S. Bose, C. E. Hong, M. E. Uddin, P. Khanra, N. H. Kim, J. H. Lee, Preparation of functionalized graphene/linear low density polyethylene composites by a solution mixing method, Carbon 49 (3) (2011) 1033–1037.

[33] S. Bose, T. Kuila, M. E. Uddin, N. H. Kim, A. K. Lau, J. H. Lee, In-situ synthesis and characterization of electrically conductive polypyrrole/graphene nanocomposites, Polymer 51 (25) (2010) 5921–5928.

[34] A. K. Geim, K. S. Novoselov, The rise of graphene, Nature materials 6 (3) (2007) 183–191.

[35] X. Li, W. Cai, J. An, S. Kim, J. Nah, D. Yang, R. Piner, A. Velamakanni, I. Jung, E. Tutuc, et al., Large-area synthesis of high-quality and uniform graphene films on copper foils, science 324 (5932) (2009) 1312–1314.

[36] K. S. Kim, Y. Zhao, H. Jang, S. Y. Lee, J. M. Kim, K. S. Kim, J.-H. Ahn, P. Kim, J.-Y. Choi, H. Hong, Large-scale pattern growth of graphene films for stretchable transparent electrodes, nature 457 (7230) (2009) 706–710.

[37] S. Bae, H. Kim, Y. Lee, X. Xu, J.-S. Park, Y. Zheng, J. Balakrishnan, T. Lei, H. Ri Kim, Y. I. Song, et al., Roll-to-roll production of 30-inch graphene films for transparent electrodes, Nature nanotechnology 5 (8) (2010) 574–578.

[38] S. M. Auerbach, K. A. Carrado, P. K. Dutta, Handbook of zeolite science and technology, CRC press, 2003.

[39] S. M. Auerbach, K. A. Carrado, P. K. Dutta, Handbook of layered materials, CRC press, 2004.

[40] J. Cejka, H. van Bekkum, A. Corma, F. Schueth, Introduction to zeolite molecular sieves, Elsevier, 2007.

[41] V. Valtchev, S. Mintova, M. Tsapatsis, Ordered porous solids: recent advances and prospects, Elsevier, 2011.

[42] J. Cejka, A. Corma, S. Zones, Zeolites and catalysis: synthesis, reactions and applications, John Wiley & Sons, 2010.

[43] S. Kulprathipanja, Zeolites in industrial separation and catalysis, John Wiley & Sons, 2010.

[44] Y. Zheng, X. Li, P. K. Dutta, Exploitation of unique properties of zeolites in the development of gas sensors, Sensors 12 (4) (2012) 5170–5194.

[45] C. Chuapradit, L. R. Wannatong, D. Chotpattananont, P. Hiamtup, A. Sirivat, J. Schwank, Polyaniline/zeolite lta composites and electrical conductivity response towards co, Polymer 46 (3) (2005) 947–953.

[46] X. Ma, H. Xu, G. Li, M. Wang, H. Chen, S. Chen, Gas-response studies of polyaniline composite film containing zeolite to chemical vapors, Macromolecular Materials and Engineering 291 (12) (2006) 1539–1546.

[47] L. Wannatong, A. Sirivat, Polypyrrole and its composites with 3a zeolite and polyamide 6 as sensors for four chemicals in lacquer thinner, Reactive and Functional Polymers 68 (12) (2008) 1646–1651.

[48] P. Phumman, S. Niamlang, A. Sirivat, Fabrication of poly (p-phenylene)/zeolite composites and their responses towards ammonia, Sensors 9 (10) (2009) 8031–8046.

[49] B. Soontornworajit, L. Wannatong, P. Hiamtup, S. Niamlang, D. Chotpattananont, A. Sirivat, J. Schwank, Induced interaction between polypyrrole and so2 via molecular sieve 13x, Materials Science and Engineering: B 136 (1) (2007) 78–86.

[50] K. Thuwachaowsoan, D. Chotpattananont, A. Sirivat, R. Rujiravanit, J. W. Schwank, Electrical conductivity responses and interactions of poly (3-thiopheneacetic acid)/zeolites l, mordenite, beta and h2, Materials Science and Engineering: B 140 (1-2) (2007) 23–30.

[51] N. Densakulprasert, L. Wannatong, D. Chotpattananont, P. Hiamtup, A. Sirivat, J. Schwank, Electrical conductivity of polyaniline/zeolite composites and synergetic interaction with co, Materials Science and Engineering: B 117 (3) (2005) 276–282.

[52] S. Chakravorty, S. Bhattacharya, A. Chatzinotas, W. Chakraborty, D. Bhattacharya, R. Gachhui, Kombucha tea fermentation: Microbial and biochemical dynamics, International journal of food microbiology 220 (2016) 63–72.

[53] H. M. Azeredo, H. Barud, C. S. Farinas, V. M. Vasconcellos, A. M. Claro, Bacterial cellulose as a raw material for food and food packaging applications, Frontiers in Sustainable Food Systems 3 (2019) 7.

[54] S.-O. Dima, D.-M. Panaitescu, C. Orban, M. Ghiurea, S.-M. Doncea, R. C. Fierascu, C. L. Nistor, E. Alexandrescu, C.-A. Nicolae, B. Tric?a, et al., Bacterial nanocellulose from side-streams of kombucha beverages production: Preparation and physical-chemical properties, Polymers 9 (8) (2017) 374.

[55] K. Watson, H. Arthur, Cell surface topography of candida and leucosporidium yeasts as revealed by scanning electron microscopy, Journal of Bacteriology 130 (1) (1977) 312–317.

[56] M. Khenfouch, U. Buttner, M. Baïtoul, M. Maaza, Synthesis and characterization of mass produced high quality few layered graphene sheets via a chemical method, Graphene 3 (2) (2014) 7–13.

[57] T. Yokoi, Characterization of zeolites by advanced sem/stem techniques, SI News 7 (2016) 17–23.

[58] M. M. Dehshibi, A. Adamatzky, Fungal Machines: Sensing and Computing with Fungi, Springer Nature Switzerland, Publication Location, 2023, Ch. Complexity of Electrical Spiking of Fungi, pp. 33–60. doi:10.1007/978-3-031-38336-6_4.

[59] M. M. Dehshibi, A. Adamatzky, Electrical activity of fungi: Spikes detection and complexity analysis, Biosystems 203 (2021) 104373. doi:10.1016/j.biosystems.2021.104373.

[60] M.-S. Abdelouahab, R. Lozi, L. Chua, Memfractance: a mathematical paradigm for circuit elements with memory, International Journal of Bifurcation and Chaos 24 (09) (2014) 1430023.

[61] S. Kvatinsky, G. Satat, N. Wald, E. G. Friedman, A. Kolodny, U. C. Weiser, Memristor-based material implication (imply) logic: Design principles and methodologies, IEEE Transactions on Very Large Scale Integration (VLSI) Systems 22 (10) (2013) 2054–2066.

[62] Q. Chen, X. Wang, H. Wan, R. Yang, A logic circuit design for perfecting memristor-based material implication, IEEE Transactions on Computer-Aided Design of Integrated Circuits and Systems 36 (2) (2016) 279–284.

[63] L. Chua, Memristor-the missing circuit element, IEEE Transactions on circuit theory 18 (5) (1971) 507–519.

[64] B. Strukov, G. S. Snider, D. R. Stewart, R. S. Williams, The missing memristor found, Nature 453 (7191) (2008) 80–83.

[65] I. Vourkas, G. C. Sirakoulis, Emerging memristor-based logic circuit design approaches: A review, IEEE circuits and systems magazine 16 (3) (2016) 15–30.

[66] X. Sun, G. Li, L. Ding, N. Yang, W. Zhang, Unipolar memristors enable “stateful” logic operations via material implication, Applied Physics Letters 99 (7) (2011).

[67] S. Kvatinsky, D. Belousov, S. Liman, G. Satat, N. Wald, E. G. Friedman, A. Kolodny, U. C. Weiser, Magic—memristor-aided logic, IEEE Transactions on Circuits and Systems II: Express Briefs 61 (11) (2014) 895–899.

[68] E. Linn, R. Rosezin, S. Tappertzhofen, U. Böttger, R. Waser, Beyond von neumann—logic operations in passive crossbar arrays alongside memory operations, Nanotechnology 23 (30) (2012) 305205.

[69] J. Borghetti, Z. Li, J. Straznicky, X. Li, D. A. Ohlberg, W. Wu, D. R. Stewart, R. S. Williams, A hybrid nanomemristor/transistor logic circuit capable of self-programming, Proceedings of the National Academy of Sciences 106 (6) (2009) 1699–1703.

[70] C. He, F. Zhuge, X. Zhou, M. Li, G. Zhou, Y. Liu, J. Wang, B. Chen, W. Su, Z. Liu, et al., Nonvolatile resistive switching in graphene oxide thin films, Applied Physics Letters 95 (23) (2009).

